# Using the basic reproduction ratio to quantify transmission and identify data gaps for epizootic haemorrhagic disease virus

**DOI:** 10.1101/2024.07.18.604050

**Authors:** Simon Gubbins

## Abstract

Epizootic haemorrhagic disease virus (EHDV) is an arbovirus transmitted by *Culicoides* biting midges that has recently emerged in Europe. Here, the basic reproduction ratio (*R*_0_) was used to quantify the transmission of EHDV and its dependence on temperature for cattle and deer. Using data from the published literature the parameters needed to calculate *R*_0_ were estimated with Bayesian methods to incorporate uncertainty in the calculations. The Sobol method of sensitivity analysis was used to determine the parameters having the greatest influence on *R*_0_ and, hence, to identify important data gaps. Depending on the strain the maximum *R*_0_ for EHDV varied from 0.7 to 2.5 in cattle and 1.3 to 4.3 in deer. The maximum *R*_0_ occurred at temperatures between 22 and 25 °C, while the lowest temperature at which *R*_0_ exceeded one was between 16 and 20 °C. The sensitivity analysis identified the threshold temperature for virus replication, the probability of transmission from host to vector and the vector to host ratio as the most important parameters influencing *R*_0_. Furthermore, there are only limited data on EHDV in European deer species and on transmission in wildlife and at the livestock/wildlife interface. These data gaps should be the focus of future research.

## 1 Introduction

Epizootic haemorrhagic disease virus (EHDV) is an arbovirus of the genus *Orbivirus*, which also includes bluetongue virus (BTV) and African horse sickness virus (AHSV). It is transmitted by *Culicoides* biting midges and can infect a wide range of wild and domestic ruminants [1,2]. It causes epizootic haemorrhagic disease (EHD), which can be particularly severe disease in wild deer, especially white-tailed deer, while clinical signs tend to be milder in cattle [3]. Historically, outbreaks of EHD have been reported in North and South America, Africa, Asia and Australasia [2,4]. In 2022 the first outbreaks of EHD were reported in Europe, caused by a strain of EHDV serotype 8. These outbreaks were detected in Sicily and Sardinia in October 2022 and in southern Spain in November 2022, and probably occurred as a result of incursions from North Africa [5]. The virus subsequently spread through the Iberian Peninsula, reaching France in 2023 [6,7].

To date limited attention has been given to quantifying the transmission of EHDV. Only one study has considered the transmission of EHDV in cattle [8], but did not integrate all the available data or consider data gaps. Here the transmission of EHDV is explored using the basic reproduction ratio, *R*_0_. This quantity is defined as the average number of secondary cases that arise from a single primary case in an otherwise susceptible population [9]. Because the introduction of an infectious disease can only result in an outbreak if *R*_0_>1, this is a means of quantifying risk, such as has been done previously for BTV [10] and AHSV [11]. Furthermore, *R*_0_ is a useful way of identifying host, virus, vector and environmental factors that influence transmission [12] and, thus, where there are important data gaps.

The approach taken in the present study was to use a previously published expression for *R*_0_ for *Culicoides*-borne viruses [10,12] to identify parameters needed to calculate *R*_0_ for EHDV. Suitable experimental and field data were extracted from the published literature and used to estimate these parameters using Bayesian methods. Once posterior distributions for each parameter had been obtained, uncertainty analysis was used to calculate *R*_0_ for EHDV. Sensitivity analysis, specifically the Sobol method [13,14], was then used to identify the most important parameters influencing the magnitude of *R*_0_ and, hence, to identify important data gaps.

The focus was on the transmission of EHDV in cattle and deer, which are the principal livestock and wildlife host species, respectively [1,2]. Fewer data are available for other susceptible species, such as sheep and goats [1,2,15], so they were not considered. Furthermore, evidence from previous outbreaks suggests that these other species play only a limited role in epidemics of EHDV [16].

## 2 Methods

### 2.1 Basic reproduction ratio for EHDV

The basic reproduction ratio, *R*_0_, for *Culicoides*-borne viruses has been derived previously [10,12]. For a single host and vector and assuming negligible disease-associated mortality (as is the case for EHDV infection in cattle [3]), it is given by,

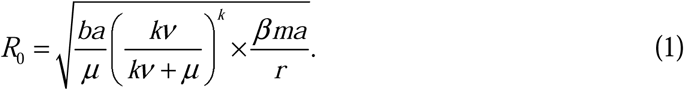

This expression for the basic reproduction ratio, (1), can be understood heuristically as follows. After a midge takes an infected blood meal, it must complete the extrinsic incubation period (EIP) before it becomes infections. Assuming that the duration of the EIP follows a gamma distribution with mean 1/ν and variance 1/*k*ν^2^ [17], the probability that the midge will survive the EIP is (*k*ν/(*k*ν+*μ*))*^k^* where *μ* is the vector mortality rate. Once a midge has completed its EIP, it will remain infectious for the rest of its lifespan, which will be 1/*μ* days on average. During this period, it will bite susceptible hosts *a* times per day (where *a* is the reciprocal of the time interval between blood meals, assumed to be equal to the biting rate), a proportion, *b*, of which will result in a newly infected host. After a host becomes infected, it will remain infectious for the duration of viraemia, which lasts 1/*r* days on average. During this time the host will be bitten by susceptible midges on average *m*×*a* times per day (here *m*=*N*/*H* is the vector to host ratio and *N* and *H* are the number of vectors and hosts, respectively) and a proportion, *β*, of these bites will result in a newly infected vector.

When there is substantial disease-associated mortality (as can be the case for EHDV infection in deer [3]), the expression for *R*_0_ given in equation (1) needs amending to account for the fact that an infected host may die before viraemia is cleared. Specifically, the mean duration for which a host remains infectious (1/*r*) is replaced by a more complex expression that accounts for this. Assuming the duration of viremia follows a gamma distribution with mean 1/*r* and variance 1/*nr*^2^ and hosts succumb to disease at a constant rate, the appropriate expression is (1/*d*)×(1-(*nr*/(*nr*+d))*^n^*) where *d* is the disease-associated mortality rate [12].

### 2.2 Parameter estimation

The parameters needed to calculate *R*_0_ for EHDV in cattle and deer were estimated from previously published data on EHDV and its *Culicoides* vectors. If suitable data for EHDV were not available, data for bluetongue virus (BTV) were used instead. The data used for parameter estimation is available in the electronic supplementary material, dataset S1 and the code to implement the methods is available online [18].

#### 2.2.1 Probability of transmission from host to vector

The probability of transmission from host to vector was estimated using data from oral infection studies using field-caught midges [19–21]. This provided the number of infected midges and the number of midges tested after feeding on a blood/virus mix via a membrane. The species tested were *C. imicola* and *C. bolitinos* [19] or *C. obsoletus* and *C. scoticus* [20,21]. Thirteen studies were included: eight using one strain of each serotype of EHDV [19]; one using a strain of EHDV-6 [20]; and four using strains of EHDV-6 and EHDV-7 [21].

The probability of transmission from host to vector for each strain, *β_s_*, was assumed to be drawn from a beta distribution with parameters *a_HV_* and *b_HV_*(i.e. *β_s_*∼Beta(*a_HV_*,*b_HV_*)). The parameters (*a_HV_* and *b_HV_*) were estimated in a Bayesian framework. The likelihood for the data is,

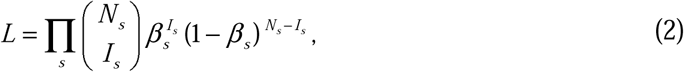

where *I_s_* is the number of infected midges, *N_s_*is the number of midges tested and *β_s_* is the probability of transmission from host to vector for strain *s*. Exponential priors with mean 100 were used for *a_HV_* and *b_HV_*. The methods were implemented in OpenBUGS (version 3.2.3; https://www.mrc-bsu.cam.ac.uk/software/). Two chains of 120,000 iterations were run, with the first 20,000 iterations discarded to allow for burn-in. The chains were then thinned (by selecting every tenth iteration) to reduce autocorrelation. Convergence was checked visually and using the Gelman-Rubin statistic implemented in OpenBUGS.

#### 2.2.2 Biting rate

The biting rate was assumed to be equal to the reciprocal of the length of the gonotrophic cycle. Data on the reciprocal of the length of the gonotrophic cycle in field-caught *C. sonorensis* (electronic supplementary material, figure S1) were extracted from the bottom panel of figure 1 in [22] using WebPlotDigitiser (version 4.6; Automeris.io) [23].

**Figure 1.**
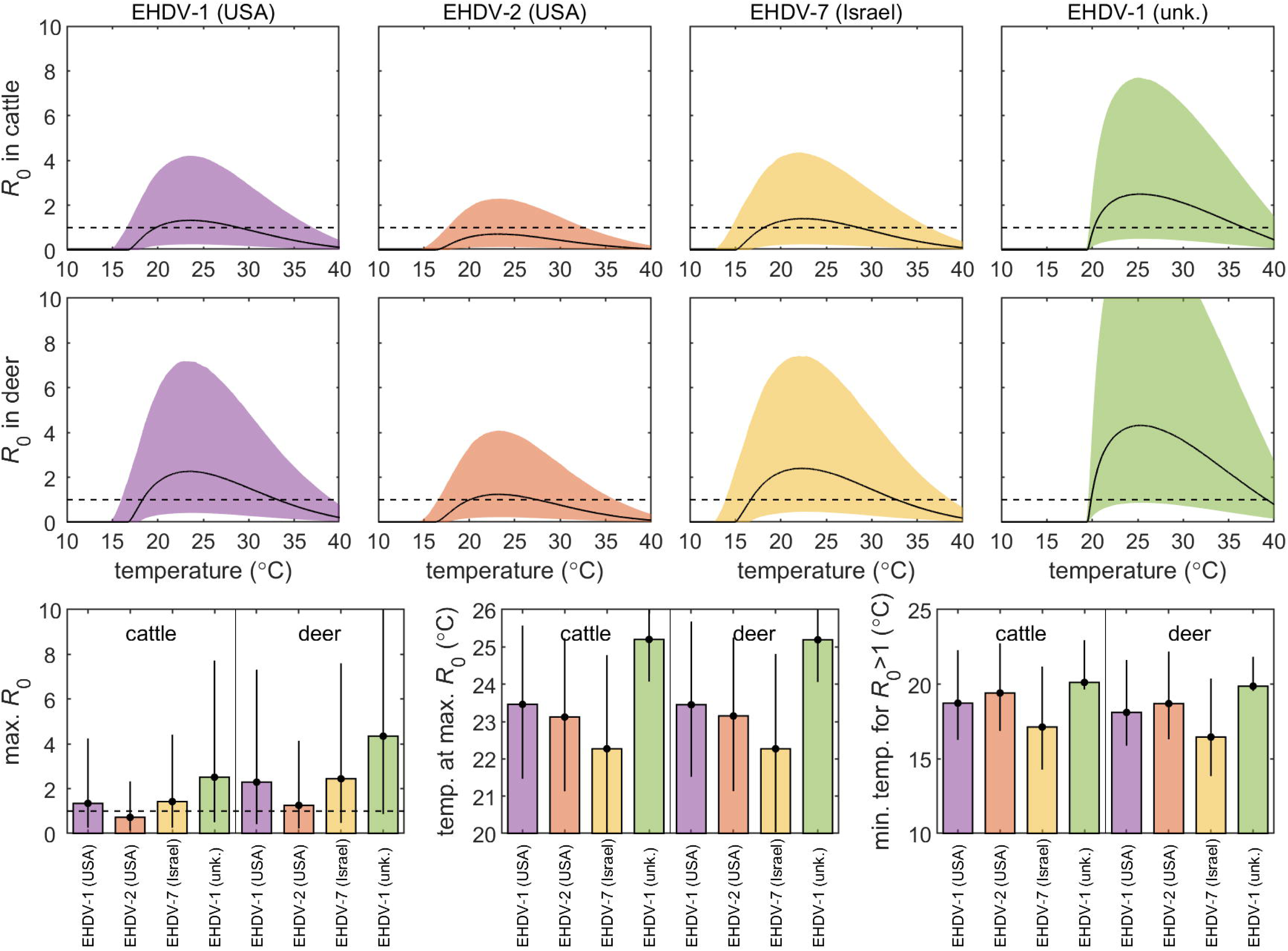
Basic reproduction ratio (*R*_0_) and its dependence on temperature for four strains of epizootic haemorrhagic disease virus in cattle or deer. The top two rows show the median (black line) and 95% prediction interval (coloured shading) for *R*_0_ as a function of environmental temperature for the strain. The bottom row shows the median (bar) and 95% prediction interval (error bars) for the maximum *R*_0_, temperature at maximum *R*_0_ and minimum temperature for *R*_0_>1 for each strain. A black dashed line indicates the threshold at *R*_0_=1. Results are based on 10,000 samples drawn from the joint posterior distribution.

The biting rate (*a*) depends on environmental temperature (*T*) and was assumed to be described by a Briere function [24], so that,

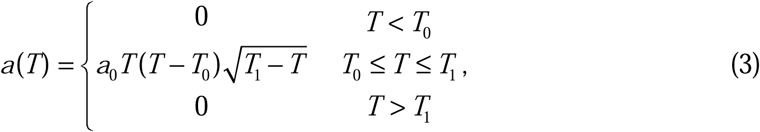

where *a*_0_ is a scale parameter and *T*_0_ and *T*_1_ are the minimum and maximum temperatures for biting, respectively. Parameters were estimated in a Bayesian framework, assuming a normal likelihood and exponential priors with mean 100 for all parameters. Because there are no data on the declining portion of equation (3) (see electronic supplementary material, figure S1), the maximum temperature for biting (*T*_1_) was fixed at 42 °C (cf. [22]). The methods were implemented using OpenBUGS. Two chains of 50,000 iterations were run, after which the chains were thinned (by selecting every fifth iteration) to reduce autocorrelation. Convergence was checked visually and using the Gelman-Rubin statistic implemented in OpenBUGS.

#### 2.2.3 Duration of viraemia in cattle

Data used to estimate the duration of viraemia in cattle were: for two calves experimentally infected with EHDV-1 or EHDV-2 [25]; for 130 cattle naturally exposed to EHDV-2, 5, 7 or 8 [1]; for four calves experimentally infected with EHDV-7 [26]; for five calves experimentally infected with EHDV-7 [27]; and for four calves experimentally infected with EHDV-8 [15]. Because of the frequency of sampling in each study, the data were used to compute the minimum and maximum duration of viraemia based on the results of virus isolation for each animal.

The duration of viraemia was assumed to follow a gamma distribution with shape parameter *n_C_* and mean 1/*r_C_*. Parameters were estimated in a Bayesian framework. The likelihood for the data was given by

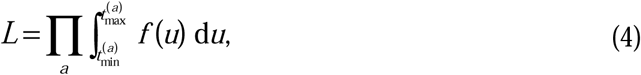

where *t*_min_^(*a*)^ and *t*_max_^(*a*)^ are the minimum and maximum duration of viraemia for animal *a*, respectively, and *f* is the probability density function for the gamma distribution. Exponential priors with mean 100 were used for all parameters.

Samples from the joint posterior density were generated using an adaptive Metropolis scheme [28], modified so that the scaling factor was tuned during burn-in to ensure an acceptance rate of between 20% and 40% for more efficient sampling of the target distribution [29]. Two chains of 120,000 iterations were run, with the first 20,000 iterations discarded to allow for burn-in of the chains. Chains were subsequently thinned (by taking every tenth sample) to reduce autocorrelation. The adaptive Metropolis scheme was implemented in Matlab (version R2020b; The Mathworks Inc.). Convergence of the scheme was assessed visually and by examining the Gelman-Rubin statistic in the coda package [30] in R (version 4.4.0) [31].

For EHDV-2 and EHDV-5 there were sufficient data to explore whether parameters (*n_C_* and 1/*r_C_*) differed between strains. Four different models were compared (electronic supplementary material, table S1) in which: (i) the shape and mean were common to both strains; (ii) the shape parameter differed between strains and the mean was common; (iii) the shape parameter was common and the mean differed; or (iv) the shape parameter and mean differed between strains. Models were compared using the deviance information criterion (DIC) [32].

#### 2.2.4 Duration of viraemia and disease-associated mortality rate in deer

Data used to estimate the duration of viraemia and disease-associated mortality rate in deer were: for 16 white-tailed deer experimentally infected with EHDV-2 [33]; for six white-tailed deer experimentally infected with EHDV-7 [34]; and for five white-tailed deer experimentally infected with EHDV-6 [35].

The duration of viraemia in deer was assumed to follow a gamma distribution with shape parameter *n_D_* and mean 1/*r_D_*. Parameters were estimated in a Bayesian framework, as described in section 2.2.3. For deer which succumbed to disease or which were still viraemic at the end of the experiment, the duration of viraemia was right-censored (i.e. the maximum duration was set to +∞).

Case fatality in deer (*f_D_*) was estimated using the same approach as described in section 2.2.2, but replacing the probability of transmission from host to vector with the case fatality and the number of infected midges and number of midges tested with the number of deer succumbing to disease and the number of deer infected, respectively. The disease-associated mortality rate (*d_D_*) can be calculated from the mean and shape for the duration of viraemia and the case fatality, using the following relationship,

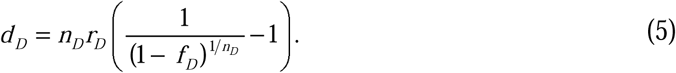

(see [11] for derivation).

#### 2.2.5 Extrinsic incubation period

The temperature dependence of the EIP was estimated from data on the infection of colonised *C. sonorensis* with different strains of EHDV [36,37]. In the experiments, *C. sonorensis* were allowed to feed on a source of virus (either on a blood/virus mix via a membrane or on an infected deer; see electronic supplementary material, table S2) and then maintained at different constant temperatures. At certain times post feeding, individual midges were tested for the level of virus present (electronic supplementary material, figure S2). A titre >2.5 log_10_ TCID_50_ [36] or >2.7 log_10_ TCID_50_ [37] was used to define a midge with a fully disseminated infection (i.e. one that is infectious).

The EIP was assumed to follow a gamma distribution with temperature-dependent mean equal to 1/ν(*T*) and variance equal to 1/*k*ν(*T*)^2^, where *k* is the shape parameter for the distribution and

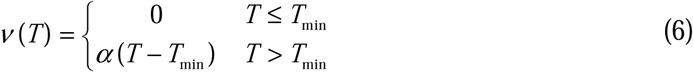

is the reciprocal of the mean EIP [17]. Here *α* is the virus replication rate and *T*_min_ is the threshold temperature (°C) for replication. This model assumes that a midge completes it EIP once it has accumulated sufficient thermal time.

The probability that a midge has a disseminated infection when tested *t* days after feeding when maintained at temperature *T* is given by,

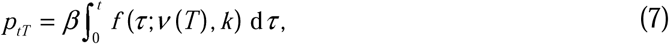

where *β* is the probability of transmission from host to vector, *f* is the probability density function for the gamma distribution, ν(*T*) is the reciprocal of the mean EIP (given by equation (6)) and *k* is the shape parameter. Differences in parameters (*β*, *T*_min_ and *α*) amongst strains and feeding routes were incorporated by allowing them to differ amongst experiments. This was incorporated by assuming hierarchical structure in the parameters such that they are drawn from higher-order distributions, so that *β*∼Beta(*a*_β_,*b*_β_), *T*_min_∼Gamma(*a_T_*,*b_T_*) and *α*∼Gamma(*a*_α_,*b*_α_), where *a_i_* and *b_i_* are the distribution parameters for the higher-order beta or gamma distributions for parameter *i*. Nine different models were compared to explore which of the parameters differed with virus strain or feeding route (electronic supplementary material, table S3).

Parameters were estimated using Bayesian methods. In this case, the likelihood for the data is given by

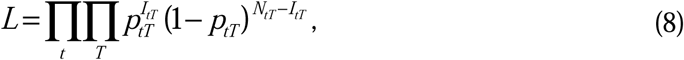

where *I_tT_* and *N_tT_* are the number of midges with a fully disseminated infection and the number of midges tested *t* days after being given an infected blood meal when maintained at temperature *T*, respectively. Hierarchical priors were used for those parameters that differed amongst strains/feeding routes and exponential priors (with mean 100) were used for the higher-order parameters in the hierarchical distributions. If a parameter was common to all strains/feeding routes, a uniform prior (with range [0,1]) was used for *β* and an exponential prior (with mean 100) was used for *T*_min_ or *α*. An exponential prior with mean 100 was used for the shape parameter (*k*).

Samples from the joint posterior density were generated using an adaptive Metropolis scheme as described in section 2.2.3. In this case, two chains of 600,000 iterations were run, with the first 100,000 iterations discarded to allow burn-in of the chains. Each chain was subsequently thinned by taking every 50th iteration.

Models assessing whether the probability of transmission from host to vector, the threshold temperature for replication or the virus replication rate differed amongst strains and feeding routes were compared using the DIC.

#### 2.2.6 Vector mortality rate

The vector mortality rate was estimated using data on the lifespan of field-caught *C. sonorensis* [38]. Data were extracted from figure 9 in [38] using WebPlotDigitiser (version 4.6; Automeris.io) [23]. The estimated lifespan was used to calculate the mortality rate (electronic supplementary material, figure S1) by assuming the mortality rate is equal to the reciprocal of the mean lifespan.

The mortality rate (*μ*) depends on environmental temperature (*T*) and the relationship can be described by

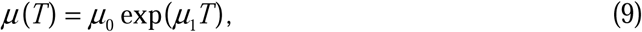

where *μ*_0_ and *μ*_1_ are parameters. These parameters were estimated in a Bayesian framework, assuming a normal likelihood and exponential priors with mean 100. The methods were implemented using OpenBUGS. Two chains of 120,000 iterations were run, with the first 20,000 iterations discarded to allow for burn-in. The chains were then thinned (by selecting every tenth iteration) to reduce autocorrelation. Convergence was checked visually and using the Gelman-Rubin statistic implemented in OpenBUGS.

#### 2.2.7 Posterior predictive checking

The fit of the models in sections 2.2.1-2.2.6 was assessed using posterior predictive checking [39]. The posterior predictive distribution was generated by simulating a single replicate of the model for each sample from the joint posterior distribution generated by the MCMC scheme. If the observed values lie within the 95% range of the posterior predictive distribution, the model was deemed to provide an adequate fit to the data.

### 2.3 Uncertainty and sensitivity analysis

To calculate *R*_0_ for EHDV in cattle or deer allowing for uncertainty in the underlying parameters, multiple sets of parameters were drawn at random from their joint posterior distributions and used to compute *R*_0_ at environmental temperatures between 10 and 40 °C.

The sensitivity of *R*_0_ to changes in each parameter was assessed using the Sobol method [13,14]. This is a variance-based global sensitivity analysis that estimates the influence of each parameter on the outputs of a model. In the context of the present study, the method quantifies the contribution of each parameter in table 1 individually and in interactions with other parameters to the total variance in *R*_0_. In particular, the first-order sensitivity index for a parameter measures the main effects of that parameter for *R*_0_ (i.e. without interactions), while the total sensitivity index measures the total effect of the that parameter (i.e. including all interactions with other parameters). When the index for a parameter is zero, *R*_0_ does not depend on that parameter, while if it is equal to one, *R*_0_ depends solely on that parameter.

**Table 1.**
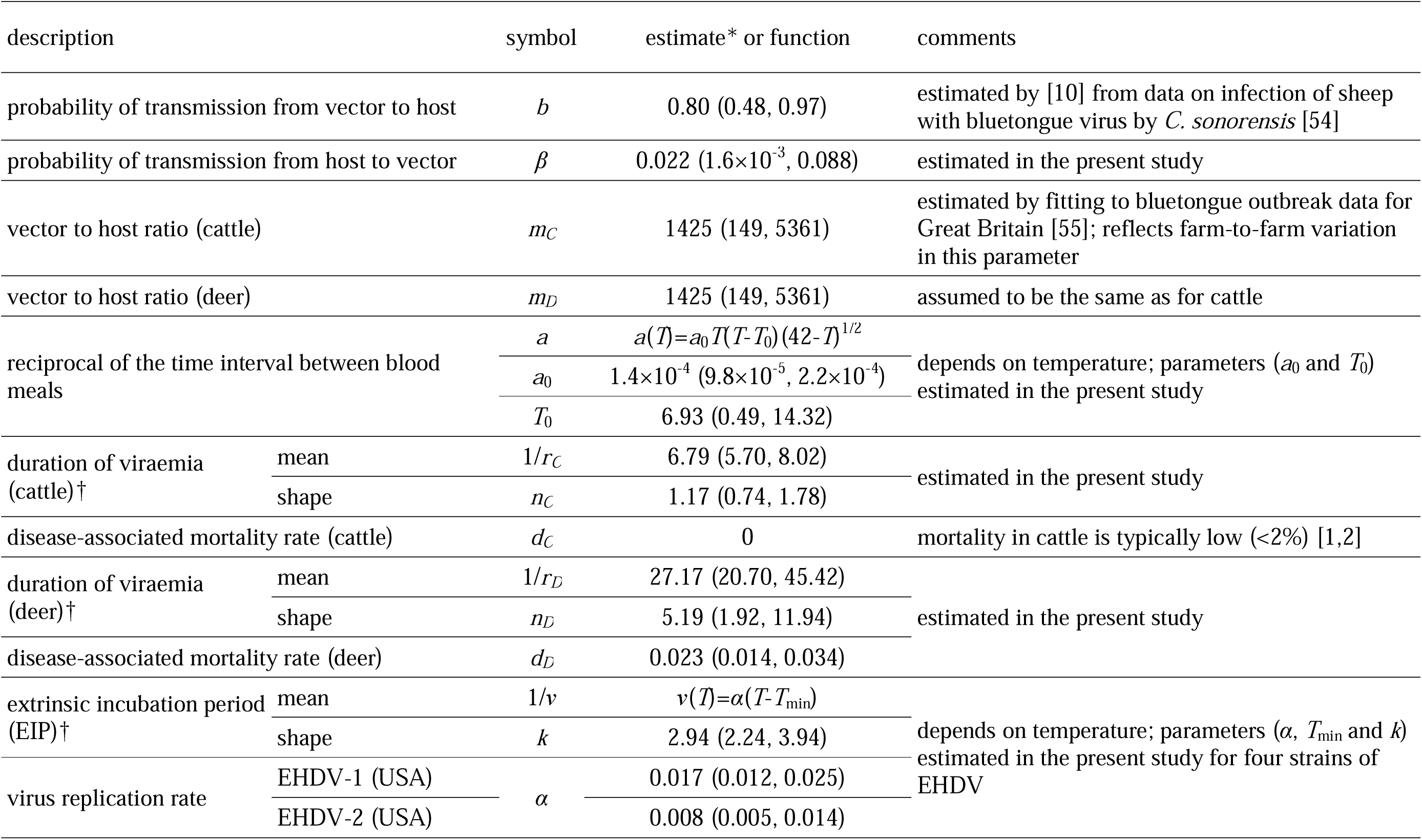
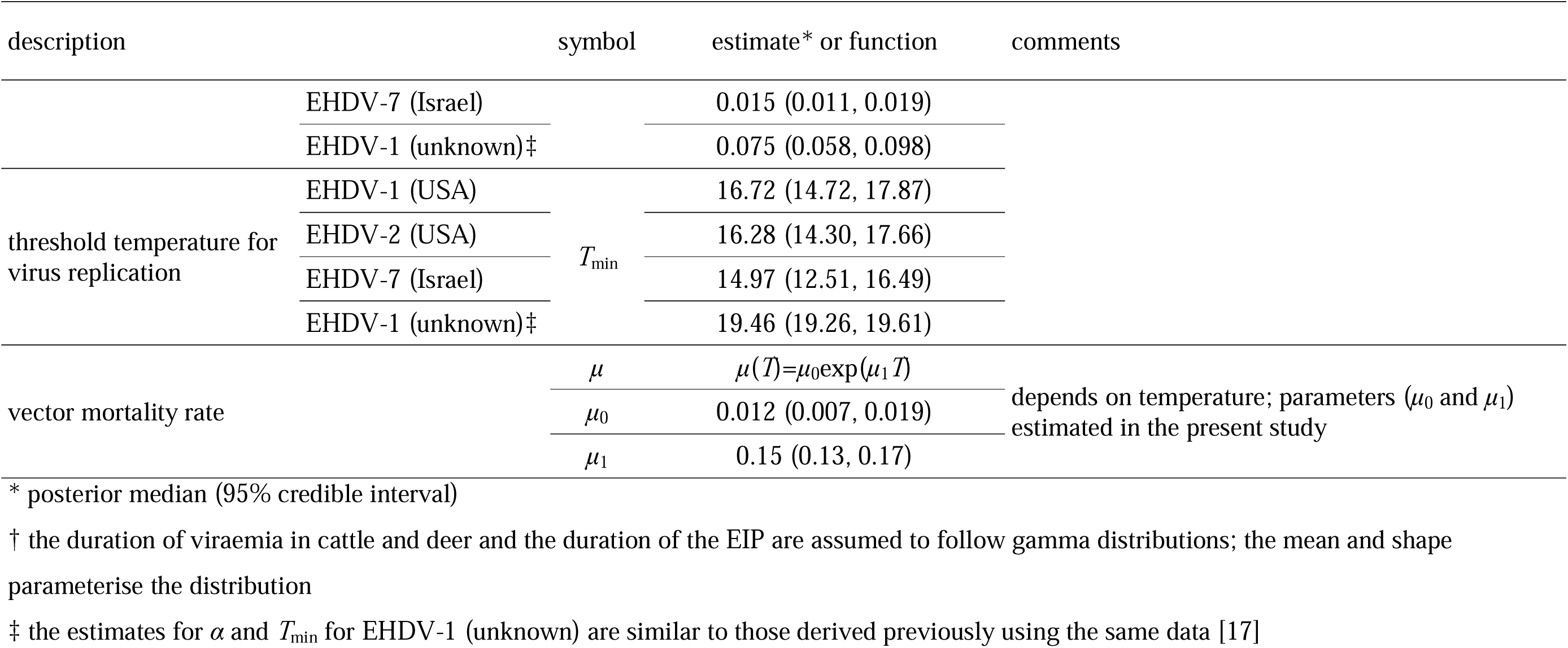
Parameters for the transmission of epizootic haemorrhagic disease virus (EHDV) in cattle and deer by *Culicoides* biting midges.

The first-order and total sensitivity indices for the parameters were calculated for *R*_0_ at different environmental temperatures using Monte Carlo methods [40]. Briefly, two random *N* by *p* matrices (where *N* is the number of samples and *p* the number of model inputs) were generated by sampling from the joint posterior distributions for the model parameters. The model (i.e. *R*_0_) was then evaluated for each set of inputs in these matrices and for combinations of the two matrices, which allows calculation of the sensitivity indices for that input (see electronic supplementary material, text S1 for details). Because they are jointly distributed (and so not independent), some parameters were grouped together as model inputs: biting rate parameters (*a*_0_, *T*_0_); host parameters (1/*r*, *n*, *d*); EIP parameters (*α*, *T*_min_, *k*); and vector mortality rate parameters (*μ*_0_, *μ*_1_). In addition, multiple replicates of the sensitivity analysis were run to check convergence of the indices.

The uncertainty and sensitivity analyses were implemented in Matlab (version R2020b; The Mathworks Inc.). The code used for the uncertainty and sensitivity analysis is available online [18].

## 3 Results

### 3.1 Parameter estimation

#### 3.1.1 Probability of transmission from host to vector

The posterior medians (95% credible interval) for *a_HV_* and *b_HV_*were 1.36 (0.59, 2.85) and 47.8 (17.6, 104.7), respectively. This gives a median probability of transmission from host to vector (*β*) of 0.022 and a 95% range of (1.6×10^-3^, 0.088) (table 1). The model adequately captured the data with the observed number of positive midges for all experiments lying within the 95% range for the posterior predictive distribution (electronic supplementary material, figure S3).

#### 3.1.2 Biting rate

The posterior medians (95% credible interval) for the slope (*a*_0_) and minimum temperature (*T*_0_) were 1.4×10^-4^ (9.8×10^-5^, 2.2×10^-4^) and 6.93 (0.49, 14.32), respectively (table 1). The fitted function, (3), and data are shown in electronic supplementary material, figure S1. The model provided an adequate fit to the data with all observed biting rates close to the median of the posterior predictive distribution (electronic supplementary material, figure S4).

#### 3.1.3 Duration of viraemia in cattle

When fitting the model to data for all strains combined the posterior medians (95% credible intervals) for the shape parameter (*n_C_*) and the mean duration (1/*r_C_*) were 1.17 (0.74, 1.78) and 6.79 (5.70, 8.02) days, respectively (table 1). The fit of the model to the data was adequate with the observed numbers of cattle with a duration of viraemia in each range lying within the 95% range for the posterior predictive distribution (electronic supplementary material, figure S5).

When fitting to data for EHDV-2 and EHDV-5 only, there was evidence that the shape parameter (*n_C_*) differed between the strains, but not the mean duration (1/*r_C_*) (electronic supplementary material, table S1). In this case, the posterior medians (95% credible intervals) for *n_C_* were 1.41 (0.79, 2.35) and 0.20 (0.05, 0.92) for EHDV-2 and EHDV-5, respectively. The posterior median (95% credible interval) for 1/*r_C_* was 7.46 (6.01, 9.13) days, which is similar to that obtained for all strains combined.

#### 3.1.4 Duration of viraemia and disease-associated mortality rate in deer

The posterior medians (95% credible intervals) for the shape parameter (*n_D_*) and mean duration (1/*r_D_*) were 5.19 (1.92, 11.94) and 27.17 (20.70, 45.42) days, respectively (table 1). The fit of the model to the data was adequate with posterior predictive *P*-values >0.05 for all but two observations (electronic supplementary material, figure S6).

When estimating the case fatality in deer (*f_D_*) the posterior medians (95% credible interval) for *a_F_* and *b_F_* were 71.0 (10.4, 252.6) and 87.1 (11.5, 300.9), respectively. This gives a median case fatality of 0.45 and a 95% prediction range of (0.37, 0.53). The model adequately captured the data with the observed numbers of deer succumbing to disease lying within the 95% range for the posterior predictive distribution (electronic supplementary material, figure S6).

Combining the estimates for the mean and shape parameter for the duration of viraemia and the case fatality using equation (5) yields a posterior median (95% credible interval) for the disease-associated mortality rate (*d_D_*) of 0.023 (0.014, 0.034) (table 1).

#### 3.1.5 Extrinsic incubation period

The best-fitting model for the EIP was one in which the threshold temperature for replication (*T*_min_) and virus replication rate (*α*) differed amongst strains, but not with feeding route (electronic supplementary material, table S3). This model adequately captured the data (electronic supplementary material, figures S2 and S7), with almost all observed numbers of midges with a fully disseminated infection lying within the 95% range of the posterior predictive distribution (electronic supplementary material, figure S7). The threshold temperature for replication ranged from 15.0 °C for EHDV-7 (Israel) to 19.5 °C for EHDV-1 (unknown) (table 1). The virus replication rate also varied amongst strains, ranging from 0.008 for EHDV-2 (USA) to 0.075 for EHDV-1 (unknown) (table 1).

Estimates for the probability of transmission from host to vector (*β*) using colony reared midges (as in the EIP studies) were markedly higher than when using field-caught midges (although of different species) (electronic supplementary material, table S4; cf. table 1). In addition, the probability of transmission from host to vector was three times higher for membrane-fed midges compared with those fed on an infected deer (electronic supplementary material, table S4).

### 3.1.6 Vector mortality rate

The posterior medians (95% credible interval) for the mortality rate parameters (*μ*_0_ and *μ*_1_) were 0.012 (0.007, 0.019) and 0.15 (0.13, 0.17), respectively (table 1). The data and fitted function, (9), are shown in electronic supplementary material, figure S1. The model provided an adequate fit to the data with all but one of the observed mortality rates lying within the 95% range for the posterior predictive distribution (electronic supplementary material, figure S4).

### 3.2 Basic reproduction ratio for EHDV

The virus replication rate and the threshold temperature for virus replication varied significantly amongst EHDV strains, though there was no evidence for strain variation amongst any of the other model parameters. Accordingly, the basic reproduction ratio (*R*_0_) was calculated separately for each of the four strains for which *α* and *T*_min_ were estimated.

There is considerable uncertainty in the predictions for *R*_0_ in cattle and sheep for all four strains (figure 1), reflecting the uncertainty in many of the underlying parameters (table 1). However, there are discernible trends in the dependence of *R*_0_ on environmental temperature and in differences in *R*_0_ amongst the strains and between host species. For all strains and host species, the basic reproduction ratio increased once the threshold temperature for virus replication was exceeded, reached a maximum level and then declined (figure 1). Comparing strains, the maximum *R*_0_ was highest for EHDV-1 (unknown) followed by EHDV-1 (USA) then EHDV-7 (Israel) and EHDV-2 (USA). The median prediction for the maximum *R*_0_ was between 0.7 and 2.5 in cattle and 1.3 and 4.3 in deer. In particular, that for EHDV-2 (USA) in cattle did not exceed the threshold at *R*_0_=1, though the upper 95% prediction limit was above one. The maximum *R*_0_ occurred at temperatures between 22.3 and 25.1 °C, with the lowest temperature for EHDV-7 (Israel) and the highest for EHDV-1 (unknown). Finally, the threshold at *R*_0_=1 was exceeded at the lowest temperature (16.5 °C) for EHDV-7 (Israel) and at the highest temperature (19.9 °C) for EHDV-1 (unknown). These patterns were the same for both cattle and deer, but *R*_0_ for all strains was higher in deer than in cattle.

### 3.3 Sensitivity analysis

The first-order and total sensitivity indices for *R*_0_ for EHDV in cattle and deer are shown in figure 2. The patterns in the sensitivity of *R*_0_ to the underlying parameters were the same for cattle and deer. Both indices indicate that the sensitivity of *R*_0_ to changes in underlying parameters depends on environmental temperature for all four strains. At lower temperatures (<18 °C) *R*_0_ is sensitive only to the EIP parameters (and the threshold temperature for virus replication, *T*_min_, in particular), and none of the others. However, at higher temperatures (>20 °C) *R*_0_ is most sensitive to the probability of transmission from host to vector (*β*) and the vector to host ratio (*m*), with these two parameters contributing approximately equally. The remaining parameters (or groups of parameters) had very limited impact on *R*_0_, either as main effects (all first-order indices <0.1) or when interactions with other parameters are also considered (all total indices <0.1).

**Figure 2.**
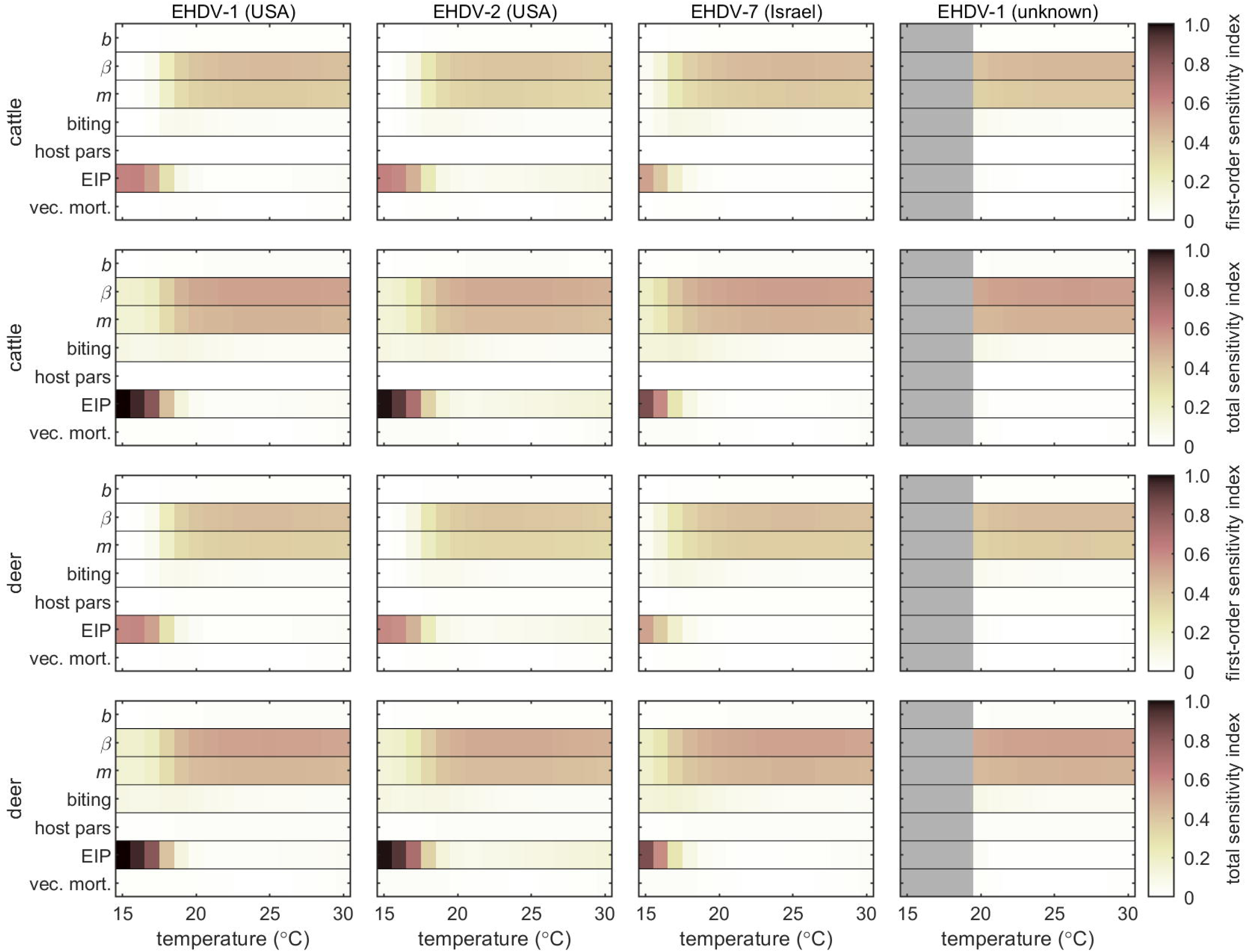
Sensitivity of the basic reproduction ratio, *R*_0_, for transmission of four strains of epizootic haemorrhagic disease virus in cattle or deer to underlying parameters and how this varies with environmental temperature. Each plot shows the first-order or total Sobol sensitivity indices (indicated by the colour bar) for each parameter or group of parameters: probability of transmission from vector to host (*b*); probability of transmission from host to vector (*β*); vector to host ratio (*m*); biting rate parameters (*a*_0_, *T*_0_); host parameters (1/*r*, *n*, *d*); extrinsic incubation period (EIP) parameters (*α*, *T*_min_, *k*); and vector mortality rate parameters (*μ*_0_, *μ*_1_). Results are the median of ten replicates with 10,000 samples drawn from the joint posterior distribution for each replicate.

## 4 Discussion

In this study the temperature-dependent basic reproduction ratio, *R*_0_, for EHDV was calculated in cattle and deer populations, the most important livestock and wildlife hosts of this virus. The parameters needed to calculate *R*_0_ were estimated from previously published data and their influence on predictions of *R*_0_ were assessed by computing Sobol sensitivity indices. Any conclusions drawn from such uncertainty and sensitivity analyses are valid only over the parameter ranges considered. However, these ranges (and the corresponding distributions) were derived from the best available data on EHDV and its *Culicoides* vectors. The only exception was the probability of transmission from vector to host, for which no data were available and was estimated based on BTV transmission to sheep instead.

The prediction intervals derived for *R*_0_ for EHDV were very wide, ranging from below the threshold at *R*_0_=1 to substantially above it (figure 1). This uncertainty reflects the variation in the underlying parameters, especially in those identified as important in the sensitivity analysis: the probability of transmission from host to vector (*β*) and the vector to host ratio (*m*) (figure 2). The variation in *β* reflects the wide range in this parameter observed for different strains and vector species [19–21]. However, the variation in *m* is more reflective of farm-to-farm variation in vector abundance (table 1) rather than uncertainty per se. Consequently, there are likely to be geographic locations where vector abundance is sufficiently high such that the lower 95% prediction interval for *R*_0_ is above one. Regardless of the uncertainty there are regions of parameter space for all four strains of EHDV considered in this study for which *R*_0_>1 at temperature relevant to much of Europe (figure 1). Consequently, it is reasonable to conclude that EHDV poses a risk to European cattle and deer. Furthermore, this is consistent with the observed spread of EHDV-8 following its introduction to southern Europe in 2022 and subsequent spread in 2023 [5–7].

The predicted values of *R*_0_ for EHDV in cattle obtained in the present study are different to those for the two other *Culicoides*-borne viruses that affect cattle and that have previously spread widely in Europe: BTV and Schmallenberg virus (SBV) (figure 3). In particular, the magnitude of *R*_0_ for EHDV (0.7-2.5) is much lower than for BTV (3.3) or SBV (4.5). Moreover, the minimum temperature for which *R*_0_>1 is lower for BTV (14 °C) and SBV (13 °C) than for EHDV (18-20 °C), as is the temperature at which *R*_0_ has its maximum (BTV: 21 °C; SBV: 21 °C; EHDV: 22-25 °C). Similar conclusions about the reduced transmission potential for EHDV in cattle (specifically, the EHDV-1 (unknown) strain) compared with BTV and SBV were obtained in another recent study [8]. However, because the authors did not calculate *R*_0_ for any of the viruses, direct comparison with the present study is not possible.

**Figure 3.**
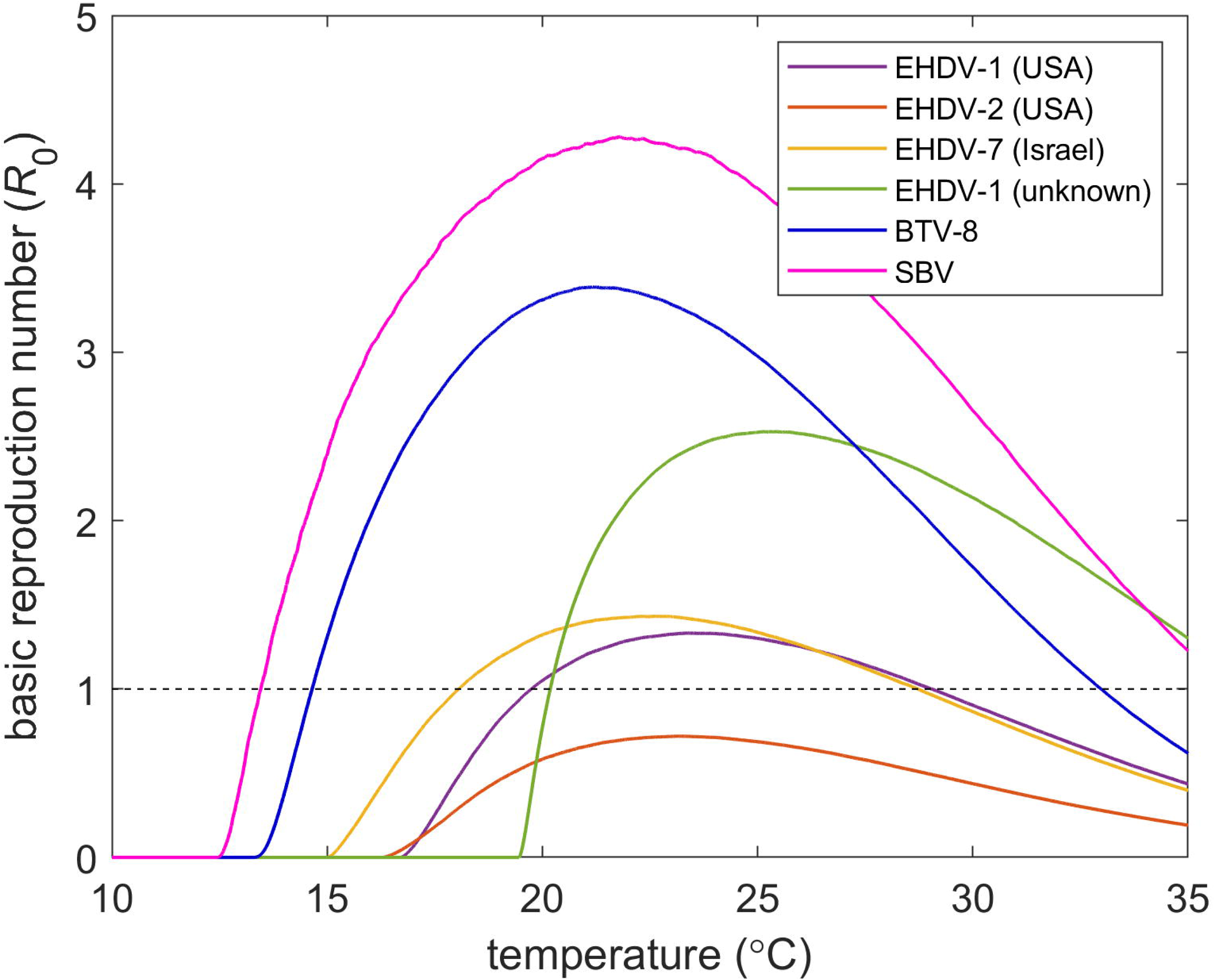
Comparison of the basic reproduction ratio (*R*_0_) in cattle for epizootic haemorrhagic disease virus (EHDV), bluetongue virus (BTV) and Schmallenberg virus (SBV). Each curve shows the posterior median for *R*_0_ and its dependence on environmental temperature for a virus/strain: EHDV-1 (USA) (purple); EHDV-2 (USA) (orange); EHDV-7 (Israel) (yellow); EHDV-1 (unknown) (green); BTV-8 (northern Europe, 2006-2010) (blue); and SBV (northern Europe, 2011) (magenta).

Although *R*_0_ only considers spread at a local spatial scale, differences in the magnitude of *R*_0_ amongst viruses would also be reflected in differences in their speed of spatial spread, with EHDV spreading more slowly that either BTV or SBV in the same region. The rate of spread has been estimated for epidemics of BTV serotype 1 in Andalusia in 2007 and in southwestern France in 2008 [41], both regions where there were outbreaks of EHDV-8 in 2022-2023. Corresponding estimates have yet to be published for EHDV-8 in either region, but comparison of maps showing spread in southwestern France suggest that the rate of spread of EHDV-8 [6,7] was slower than for BTV-1 [41]. This is consistent with a lower *R*_0_ for EHDV compared with BTV, but perhaps not one so low as predicted in the present study. For example, the difference in *R*_0_ between BTV (3.3) and SBV (4.5) was sufficient to explain the considerably greater spread of SBV compared with BTV in northern Europe [12]. These differences in *R*_0_ were a consequence of higher vector competence and faster replication within vectors at lower temperatures for SBV compared with BTV [12].

The magnitude of *R*_0_ for EHDV was highly sensitive to both the probability of transmission from host to vector (*β*; i.e. vector competence) and the EIP parameters (figure 2). Vector competence has not yet been reported for the strain of EHDV-8 currently circulating in Europe for any *Culicoides* species, though it has been detected in pools of *C. obsoletus/scoticus*, *C. imicola* and *C. punctatus* collected in Sardinia [42]. The minimum prevalence for *C. obsoletus/scoticus* (i.e. no. positive pools/total no. insects in the pools) was 0.11% (4/3542). This is comparable to the minimum prevalence for SBV (0.14%; 2/1440) [43] and higher than for BTV-8 (0.05%; 1/2200) [44]. This suggests that vector competence for EHDV-8 may be similar to that for SBV and higher than that for BTV-8. It would also indicate that the vector competence is likely to be at the higher end of the distribution estimated in the present study (table 1).

Experimental work to better quantify vector competence in European *Culicoides* species should be a priority, especially for the strain of EHDV-8 currently circulating in Europe. However, analysis in the present study indicates that the probability of a vector becoming infected depends not only on EHDV strain, but also on feeding route (feeding via membrane or on an infected animal), vector species and field-caught compared with colony midges (table 1; electronic supplementary material, table S4). Other experimental work has also shown a dependence of the probability of infection on viral titre in the blood meal on which an insect fed [45–47]. Consequently, these factors should also be taken into account in the design any vector competence experiments.

The threshold temperature for virus replication (*T*_min_) determines the temperature at which *R*_0_ changes from zero to greater than zero and so constrains the transmission season for EHDV. The threshold temperatures (15-19 °C) estimated in the present study for four strains of EHDV are higher than estimated for BTV (11-14 °C) [16] or SBV (12 °C) [11]. This suggests that the transmission season could be shorter for EHDV compared with *Culicoides*-borne viruses that have previously spread in Europe, especially at more northerly latitudes. Moreover, the EIP parameters in general, and the threshold temperature in particular, are important for determining the magnitude of *R*_0_ for EHDV at temperatures close to the threshold (figure 2). Accordingly, the EIP for EHDV-8 and its dependence on temperature, which has not been quantified, is an important data gap.

A third parameter to which *R*_0_ for EHDV is sensitive is the vector to host ratio (*m*) (figure 2). The assumed variation in this parameter for cattle reflects farm-to-farm variation in midge abundance (table 1). Although various *Culicoides* species are known to feed on deer [48,49], little else is known about their association with deer populations [50], including the relationship between abundance and deer population size. In the absence of this information, the vector to host ratio was assumed to follow the same distribution as for cattle. Similarly, the biting rate on deer was assumed to be the same as that for cattle. Better characterisation of the relationships between deer and *Culicoides* biting midges would allow a more robust assessment of the role of deer in the transmission of EHDV in Europe. Evidence from previous epidemics of BTV suggest that deer played an important role in maintaining the virus and vector populations in areas of southern Europe [50]. By contrast, deer were less important in northern and central Europe, where they did not act as maintenance hosts for BTV [51].

For all four strains of EHDV considered in the present study *R*_0_ was higher in deer than in cattle (figure 1), suggesting deer could play an important role in the transmission of EHDV. However, the parameters for deer used in the present study were estimated from data on white-tailed deer infected with strains of EHDV circulating in the USA. EHDV has been widely studied in white-tailed deer [2], reflecting the impact of EHD on populations of this species in endemic areas [3]. Only limited data are available on EHDV infection in European deer species [52,53]. These demonstrate that red, roe and fallow deer and muntjac are susceptible to EHDV infection and, in the case of red deer, can develop clinical disease, but are not sufficient to parameterise transmission models.

## 5 Conclusions

Results of the present study show that *R*_0_ for EHDV depends on strain, but that *R*_0_ exceeds one (and so can cause outbreaks) at temperatures relevant to much of Europe. Sensitivity analysis identified the probability of transmission from host to vector (i.e. vector competence), the threshold temperature for virus replication and the vector to host ratio as the most important parameters influencing *R*_0_. In addition, there are only limited data on EHDV in European deer species and on *Culicoides* populations at the deer/livestock interface. These areas should be the focus of future research.

## Supporting information

Previously published data used in analyses

## Data sharing

No new data were generated by this work. The data extracted from the published literature and used to parameterise the model are provided in the electronic supplementary material, dataset S1. All code used to estimate parameters and implement the uncertainty and sensitivity analyses is available online [18].

## Ethics statement

No ethical issues were raised by this work.

## Author contributions

Conceptualization: SG. Formal analysis: SG. Methodology: SG. Software: SG. Writing - original draft: SG. Writing - review & editing: SG.

## Acknowledgements

The author is grateful to Christopher Sanders and Marion England at The Pirbright Institute for helpful discussions on this work.

