## Supplementary text, tables and figures for "Using the basic reproduction ratio to quantify transmission and identify data gaps for epizootic haemorrhagic disease virus"

**Electronic supplementary material**

Simon Gubbins*

*The Pirbright Institute, Ash Road, Pirbright, Surrey GU24 0NF, UK*

**This PDF contains:**

Text S1

Figures S1-S7

Tables S1-S4

**Additional supplementary files are:**

**Dataset S1.** Data extracted from the published literature and used to parameterise the basic reproduction ratio for EHDV.

**Text S1 Monte Carlo methods for computing Sobol sensitivity indices**

The first-order ($s_i$) and total ($s_{Ti}$) sensitivity indices for input $i$ (i.e. parameter or group of parameters) were calculated for different temperatures using Monte Carlo methods [1]. Two random $N$ by $p$ matrices (where $N$ is the number of samples and $p$ the number of model inputs), **A** and **B**, were generated by sampling from the joint posterior distributions for the model parameters. For each input (i.e. parameter or group of parameters), two further matrices were generating combining elements of **A** and **B**, specifically $\mathbf{A}_B^{(i)}$ which is matrix **A** except for the columns relating to input $i$ which come from matrix **B** and $\mathbf{B}_A^{(i)}$ which is matrix **B** except for the columns relating to input $i$ which come from matrix **A**. For each row $r$ of these four matrices, the basic reproduction ratio ($R_0$) was computed at each temperature, and the values obtained were used to the calculate the sensitivity indices, $s_i$ and $s_{Ti}$, for input $i$ as follows:

$$s_i = 2 \times \frac{\sum_{r=1}^{N} (R_0(\mathbf{B}_A^{(i)})_r - R_0(\mathbf{B})_r)(R_0(\mathbf{A})_r - R_0(\mathbf{A}_B^{(i)})_r)}{\sum_{r=1}^{N} \left[ (R_0(\mathbf{A})_r - R_0(\mathbf{B})_r)^2 + (R_0(\mathbf{B}_A^{(i)})_r - R_0(\mathbf{A}_B^{(i)})_r)^2 \right]},$$

and

$$s_{Ti} = \frac{\sum_{r=1}^{N} \left[ (R_0(\mathbf{B})_r - R_0(\mathbf{B}_A^{(i)})_r)^2 + (R_0(\mathbf{A})_r - R_0(\mathbf{A}_B^{(i)})_r)^2 \right]}{\sum_{r=1}^{N} \left[ (R_0(\mathbf{A})_r - R_0(\mathbf{B})_r)^2 + (R_0(\mathbf{B}_A^{(i)})_r - R_0(\mathbf{A}_B^{(i)})_r)^2 \right]}.$$

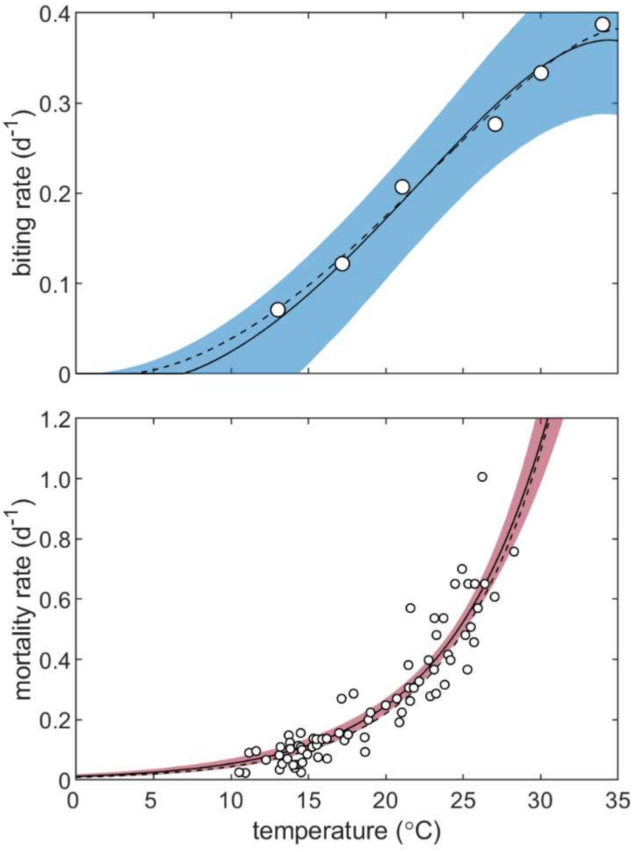


**Figure S1.** Life history parameters for *Culicoides* biting midges and their dependence on environmental temperature: reciprocal of the time interval between blood meals (extracted from [2]), assumed to be equal to the biting rate (top panel), and vector mortality rate (extracted from [3]) (bottom panel). Each plot shows the observed values (circles) extracted from previous published literature [2,3], posterior median (black line) and 95% credible interval (shading) for the parameter. The black dashed lines show the previously published curves estimated by Mullens et al. [2] and Gerry & Mullens [3], respectively.

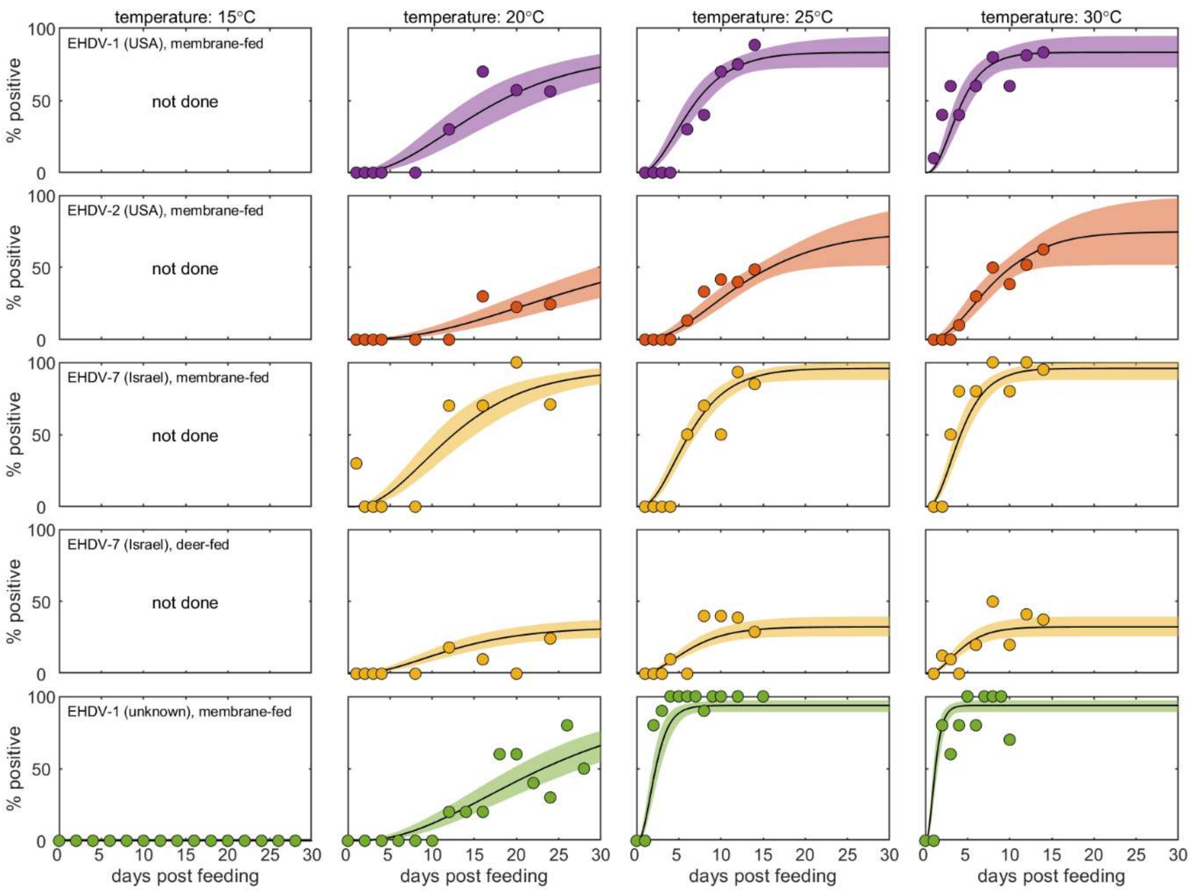


**Figure S2.** Proportion of *Culicoides* biting midges with a fully disseminated infection when infected with four strains of epizootic haemorrhagic disease virus (EHDV) by different feeding routes (rows) and maintained at different temperatures (columns). Each plot shows the observed values (circles) extracted from the previously published literature [4,5], posterior median (black line) and 95% credible interval (shading) for the proportion of midges with a fully disseminated infection.

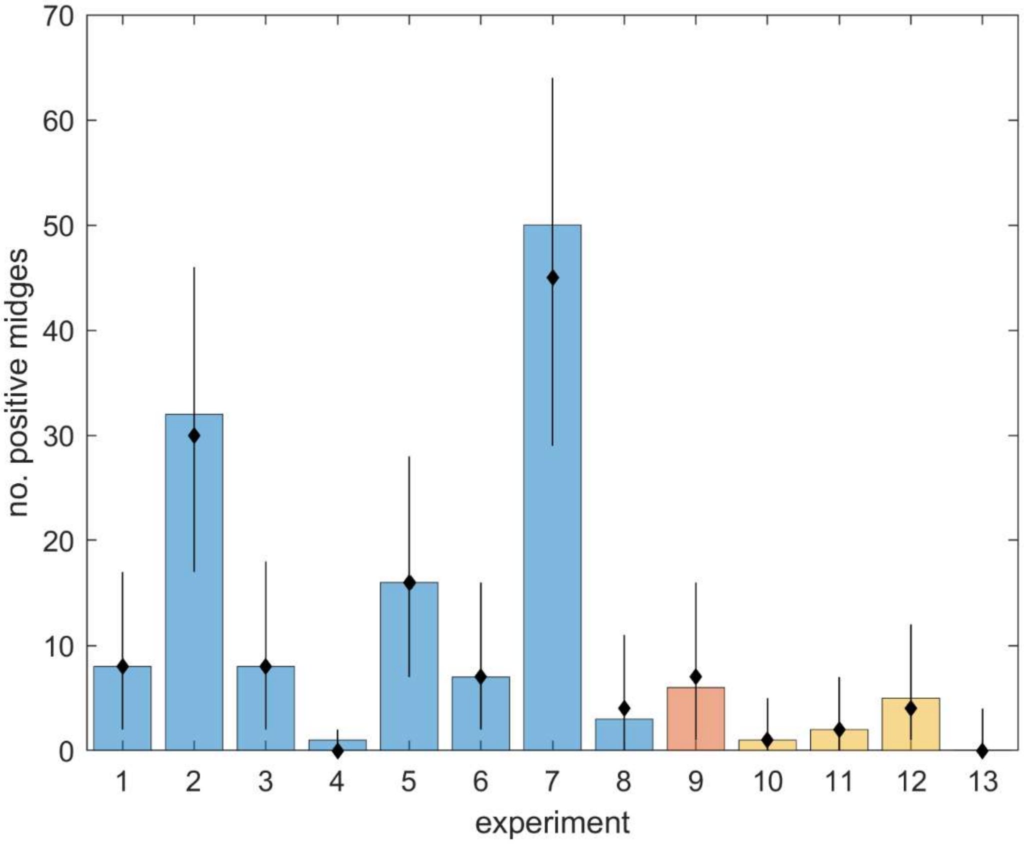


**Figure S3.** Posterior predictive checking of the model for the probability of transmission from host to vector for epizootic haemorrhagic disease virus (EHDV) in *Culicoides* biting midges. Bars show the observed number of positive midges from previously published literature (blue, [6]; orange, [7]; yellow, [8]) and black diamonds and error bars show the median and 95% range, respectively, for the posterior predictive distribution.

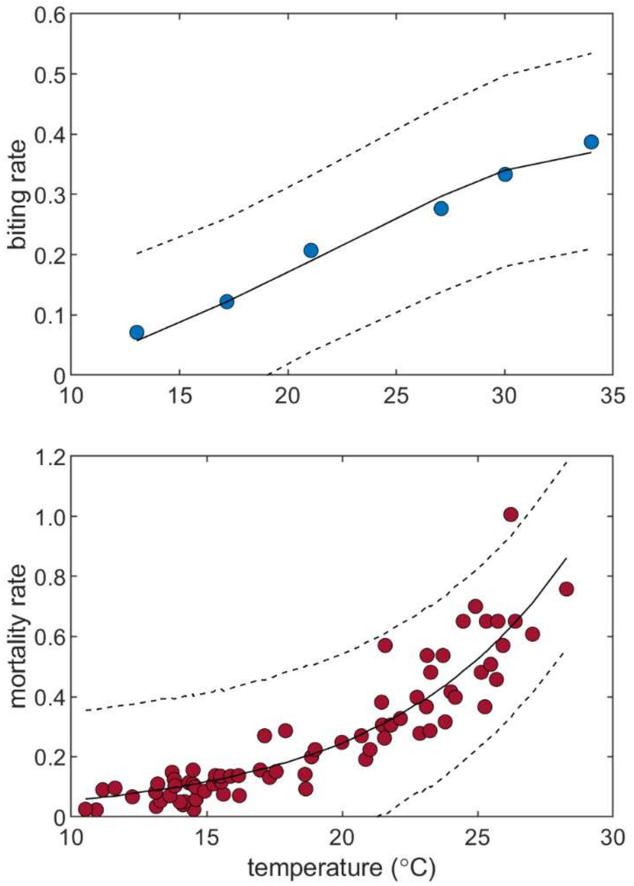


**Figure S4.** Posterior predictive checking of the temperature dependent models for the reciprocal of the time interval between blood meals, assumed to be equal to the biting rate (top panel), and the vector mortality rate (bottom panel). Each plot shows the observed values (circles) extracted from previous published literature [2,3] and the median (solid line) and 95% range (dashed lines) for the posterior predictive distribution.

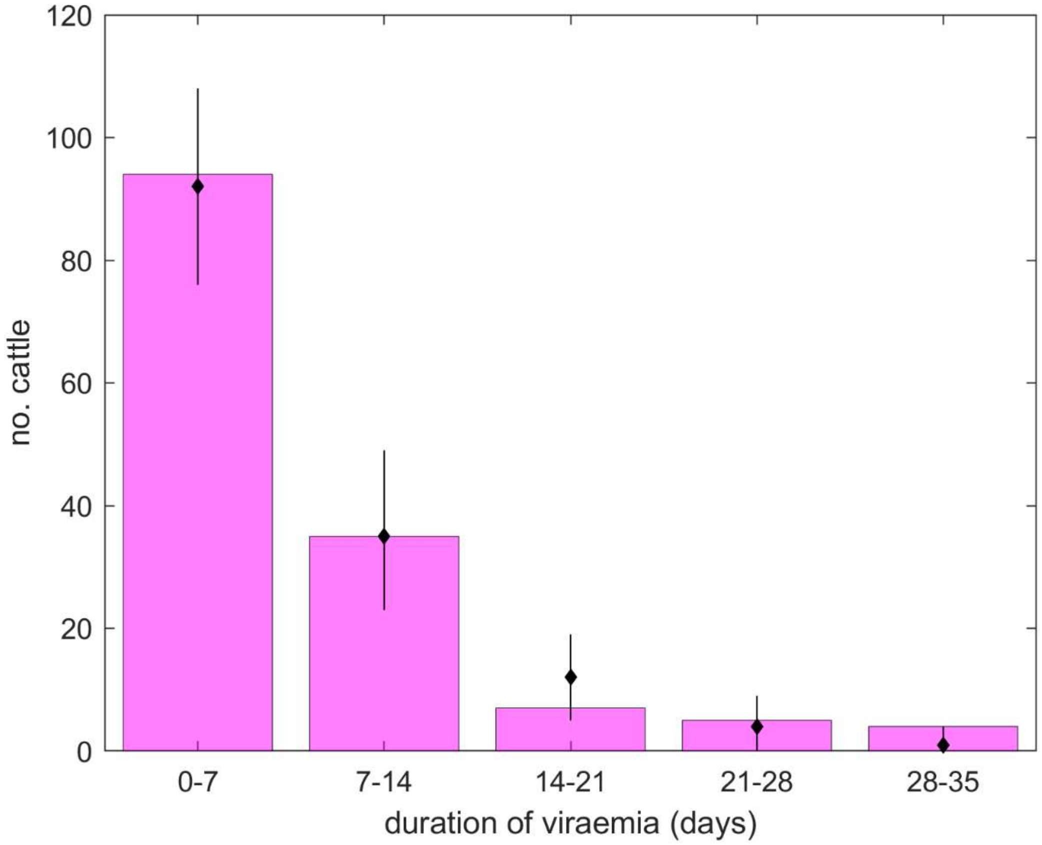


**Figure S5.** Posterior predictive checking of the model for the duration of viraemia for epizootic haemorrhagic disease virus (EHDV) in cattle. Bars show the observed number of cattle with a duration of viraemia in the given range extracted from five papers [9-13] and black diamonds and error bars show the median and 95% range, respectively, for the posterior predictive distribution.

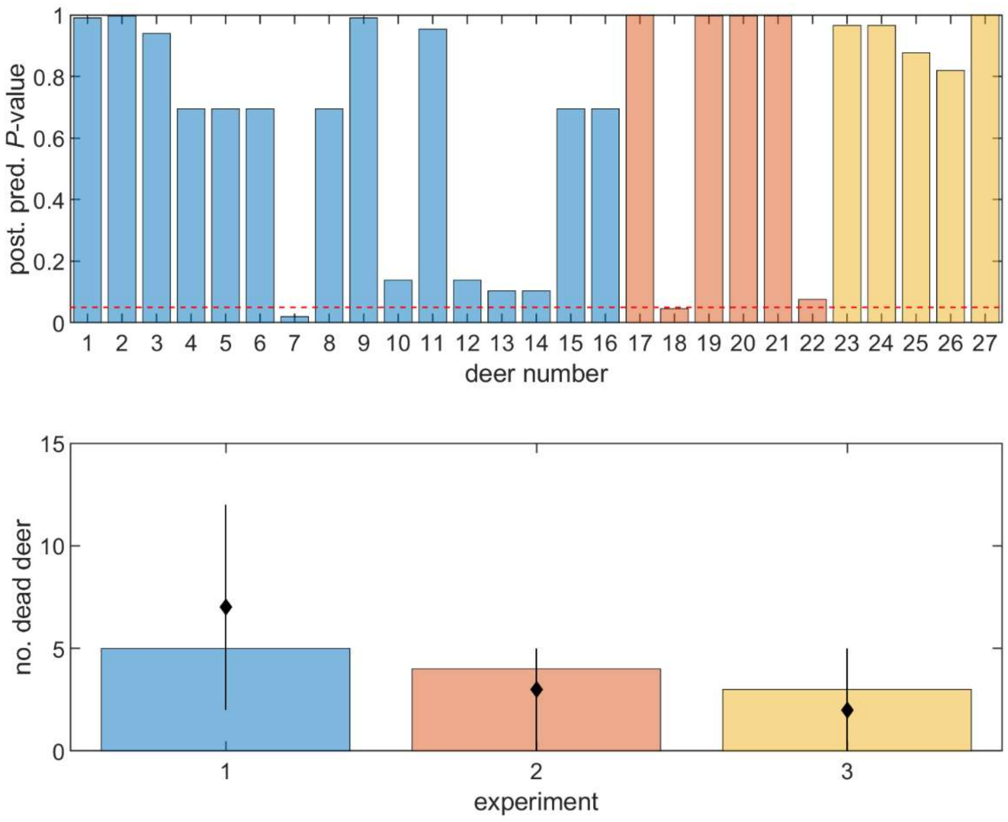


**Figure S6.** Posterior predictive checking of the model for duration of viraemia for epizootic haemorrhagic disease virus (EHDV) in deer. The top panel shows the posterior predictive *P*-values (bars) for the duration of viraemia for each animal. The red dashed line indicates the threshold at *P*=0.05. The bottom panel shows the observed number of deer succumbing to disease extracted from the previously published literature (bars) and the median (black diamonds) and 95% range (error bars) for the posterior predictive distribution. In both panels bar colour indicates the study: blue, [14]; orange, [15]; and yellow, [16].

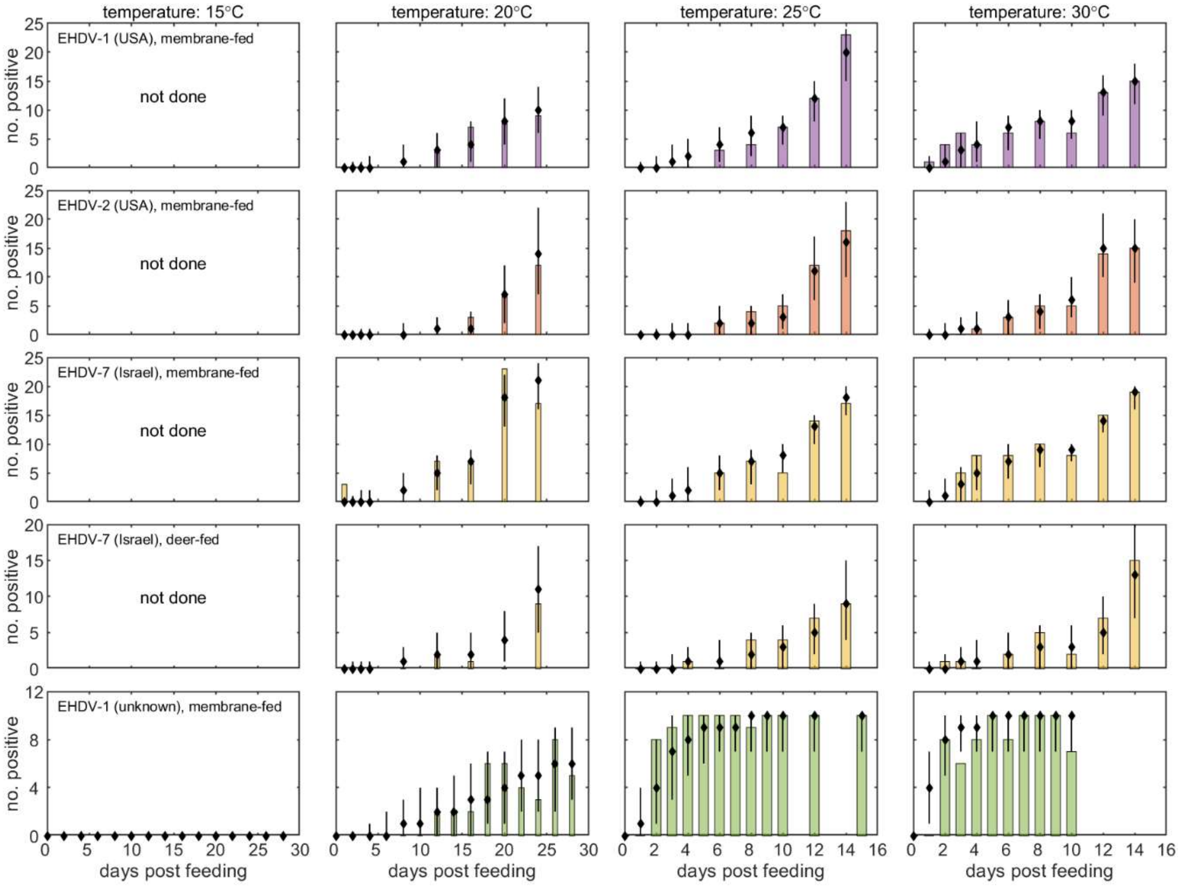


**Figure S7.** Posterior predictive checking of the model for the number of *Culicoides* biting midges with a fully disseminated infection when infected with four strains of epizootic haemorrhagic disease virus (EHDV) by different feeding routes (rows) and maintained at different temperatures (columns). Each plot shows the observed number of midges with a fully disseminated infection (bars) extracted from the previously published literature [4,5] and the median (black diamonds) and 95% range (error bars) for the posterior predictive distribution.

**Table S1.** Comparison of models* for the duration of viraemia in cattle.

| shape parameter ($n_C$) | mean duration ($1/r_C$) | DIC† |
|---|---|---|
| common | common | 234.9 |
| varies | common | **228.5**‡ |
| common | varies | 232.1 |
| varies | varies | 226.9 |

* common: parameter common to both strains; varies: parameters vary between strains

† DIC: deviance information criterion; a model with a smaller DIC is preferred to one with a higher DIC

‡ because the change in DIC between this model and the more complex one was less than 2, the simpler model was preferred on grounds of model parsimony

**Table S2.** Summary of the experimental data used to estimate the extrinsic incubation period of epizootic haemorrhagic disease virus (EHDV).

| virus* | feeding route | reference |
| --- | --- | --- |
| 1 (white-tailed deer, USA, 2006) | membrane, deer blood/virus mix | [4] |
| 2 (white-tailed deer, USA, 1993) | membrane, deer blood/virus mix | |
| 7 (Holstein cow, Israel, 2006) | membrane, deer blood/virus mix | |
| 7 (Holstein cow, Israel, 2006) | infected deer | |
| 1 (unknown†) | membrane, horse blood/virus mix | [5] |

* serotype (species, country, year isolated)

† the virus was obtained from Onderstepoort Veterinary Institute, South Africa, but details of the species, country and year isolated were not provided; however, it is likely to be distinct from the strain of EHDV-1 used in Ruder et al. [15] which was collected in 2006

**Table S3.** Comparison of models* for the replication of epizootic haemorrhagic disease virus in *Culicoides* biting midges.

| probability of transmission from host to vector (β) | threshold temperature ($T_{min}$) | viral replication rate (α) | DIC† |
|---|---|---|---|
| common | common | common | 902.6 |
| varies | common | common | 592.6 |
| common | varies | common | 768.6 |
| common | common | varies | 586.1 |
| varies | varies | common | 577.0 |
| varies | common | varies | 570.7 |
| common | varies | varies | 474.6 |
| varies | varies | varies | 465.5 |
| varies | varies‡ | varies‡ | **467.4**⁋ |

* common: parameter common to all experiments; varies: parameters vary amongst experiments (i.e. strain and feeding route)

† DIC: deviance information criterion; a model with a smaller DIC is preferred to one with a higher DIC

‡ for this model $T_{min}$ and $\alpha$ varied only amongst viral strains, but not between feeding routes

⁋ because the change in DIC between this model and the more complex one was less than 2, the simpler model was preferred on grounds of model parsimony

**Table S4.** Probability of transmission from host to vector estimated from experiments measuring the extrinsic incubation period of epizootic haemorrhagic disease virus (EHDV).

| virus and feeding route | posterior median | 95% credible limit | |
|---|---|---|---|
| | | lower | upper |
| EHDV-1 (USA), membrane fed | 0.83 | 0.73 | 0.95 |
| EHDV-2 (USA), membrane fed | 0.75 | 0.52 | 1.00 |
| EHDV-7 (Israel), membrane fed | 0.96 | 0.88 | 1.00 |
| EHDV-7 (Israel), fed on infected deer | 0.32 | 0.26 | 0.40 |
| EHDV-1 (unknown), membrane fed | 0.94 | 0.89 | 0.97 |
